# A framework for generating interactive reports for cancer genome analysis

**DOI:** 10.1101/194035

**Authors:** Ai Okada, Kenichi Chiba, Hiroko Tanaka, Satoru Miyano, Yuichi Shiraishi

## Abstract

**Summary:** We introduce paplot, the software for generating dynamic reports that are frequently necessary in the post analytical phases of cancer genome studies. The “interactive” nature of the paplot-generated reports enables users to extract much richer information than that obtained from static graphs via most conventional visualization tools.

**Availability and implementation:** The python implementation for paplot (MIT license) is available at https://github.com/Genomon-Project/paplot. The documentation is at http://paplot-doc.readthedocs.io/en/latest/.

## Introduction

The worldwide rapid accumulation of massive cancer genome sequencing data presents an unprecedented opportunity for obtaining enormous biologically and clinically important information. Several analytical tools have been developed for identifying somatic mutations (Koboldt, et al., 2012; Shiraishi, et al., 2013), structural variations (Rausch, et al., 2012), and prominent mutation signatures (Shiraishi, et al., 2015) and so on. However, the subsequent visualization and interpretation often demand us additional laborious efforts such as formatting and organizing the result tables via additional scripting, as well as graphic manipulation via drawing tools, occasionally with multiple trial-and-error parameter adjustments. Additionally, as many existing tools only provide static figures (Skidmore, et al., 2016; Yin, et al., 2012), it is necessary to again repeatedly execute drawing tools with different settings even for generating slightly different visualizations (e.g., sorting and filtering by specific criteria). Furthermore, even when some interesting characteristics are observed, we often need to bother to refer back to the raw analytical result tables for identifying the samples and the exact values of interest, as these may not be shown in the static figures.

Here, we present paplot (https://github.com/Genomon-Project/paplot), a suite of programs to generate various reports for commonly needed tasks in cancer genome analysis. paplot accepts text-based input data and configuration files and generates HTML files with dynamic, interactive data visualizations powered by D3.js (https://d3js.org). By modifying settings in the configuration files, users can freely change various visual aspects such as colors of objects and contents of pop-up windows appearing after DOM events such as mouse-click and mouse-over.

## Quality Control Report

Confirming the quality of sequence data (e.g., coverage, insert size and mapping ratio) is a fundamental step in sequence analysis. In many cases, the quality control check is performed by visual inspection of the barplot to check whether there are some samples with strangely heights. However, as adding the X-axis labels for all the samples is not always possible especially when the sample size in the cohort is large, we have to spend excessive time to retrieve the raw table data to identify the samples with the strange features. By using the Quality Control Report generated by paplot (Figure 1A), one can immediately identify such samples via pop-up contents.

**Figure 1.**
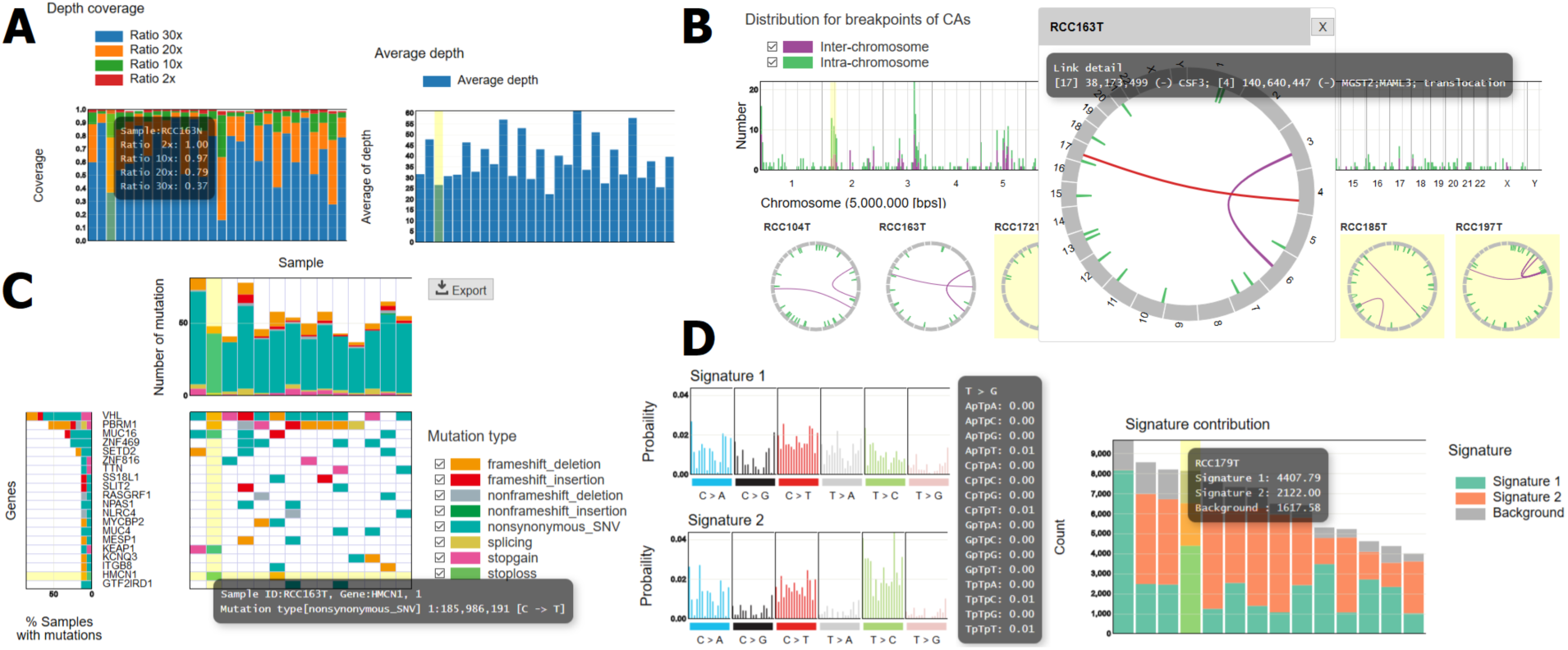
Example of interactive reports created by paplot. **(A)** Quality Control Report, **(B)** Chromosomal Aberration Report, **(C)** Mutation Matrix Report, **(D)** Mutational Signature Report.

## Chromosomal Aberration Report

Chromosomal aberrations play important roles in cancer development (Mitelman, et al., 2007). After characterizing chromosomal aberrations (genomic structural variations from whole genome sequencing data or fusion transcripts from transcriptome sequencing data), it is common to generate a visual overview of chromosomal aberrations, typically using the circos plot (Krzywinski, et al., 2009), to examine for certain specific phenomena such as chromothripsis (Stephens, et al., 2011). When the users open the Chromosomal Aberrations Report (Figure 1B), they will first find thumbnails for chromosomal aberrations across all the samples, thus providing a cohort-level landscape. On clicking one of the thumbnails, the bigger picture of chromosomal aberrations for the sample of interest appears, and detailed information of each variant (e.g., exact coordinates of breakpoints and affected genes) pops up upon mouse-over. Furthermore, the distribution for the cumulative count of breakpoints of chromosomal aberrations partitioned by a certain chromosomal bin size (default size is 50 Kbp) is automatically generated, facilitating identification of common cancer-driving variants represented by prominent peaks (Kataoka, et al., 2016).

## Mutation Matrix Report

Obtaining a comprehensive vision of somatic mutations across cancer genomes in a cohort of interest to examine frequencies of affected genes, relative ratio of variant types (e.g., missense, nonsense), and co-occurrence and exclusivity between two genes, is a principal goal in cancer genome analysis. paplot produces the Mutation Matrix Report (Figure 1C), a dynamic report illustrating the mutation status, which can fulfill the above purpose. Users can enjoy sorting by several criteria such as sample-wise and gene-wise frequencies. In addition, the user can generate a “waterfall” plot (Skidmore, et al., 2016), where samples are hierarchically divided per the mutation status of genes in descending order of mutation frequencies. Furthermore, clinical information can be included in the plot footer, which can work as additional sort keys.

## Mutational Signature Report

In the past few years, massive genome cancer sequencing analysis revealed many characteristic patterns of somatic mutations. The profile of such mutation signature is typically represented by a bar plot with 96 possible types (6 types of substitution × 4 types of 5’ base × 4 types of 3’ base) (Alexandrov, et al., 2013). We have recently proposed another way of visualizing mutation signatures to enable the inclusion of more contextual factors (Shiraishi, et al., 2015). paplot can generate the Mutational Signature Report (Figure 1D) in both format types. Furthermore, since the barplot of membership parameters (estimated fractions of operative mutation signatures for each sample) is also generated, the user can easily spot the sample with prominent mutation signatures via pop-up windows.

## Conclusion

Interactive graphics contain richer information than static ones, thus facilitating effective and intuitive interpretation of analytical results. Owing to the advancement in information technologies and development of application frameworks for easy generation of interactive graphics, using interactive graphics for academic research and auto-generated reports for clinical sequencing will become more common. We believe that paplot will be useful for academic research and clinical practice.

## Acknowledgements

The supercomputing resources were provided by Human Genome Center, The Institute of Medical Science, The University of Tokyo.

